# 3D-Cardiomics: A spatial transcriptional atlas of the mammalian heart

**DOI:** 10.1101/792002

**Authors:** Monika Mohenska, Nathalia M. Tan, Alex Tokolyi, Milena B. Furtado, Mauro W. Costa, Andrew J. Perry, Jessica Hatwell-Humble, Karel van Duijvenboden, Hieu T. Nim, Susan K. Nilsson, David R. Powell, Nadia A. Rosenthal, Fernando J. Rossello, Mirana Ramialison, Jose M. Polo

**Affiliations:** Department of Anatomy and Developmental Biology, Monash University, Wellington Road, Clayton, Victoria, Australia; Development and Stem Cells Program, Monash Biomedicine Discovery Institute, Wellington Road, Clayton, Victoria, Australia; Australian Regenerative Medicine Institute, Monash University, Wellington Road, Clayton, Victoria, Australia; The Jackson Laboratory, Bar Harbor, ME, USA; Monash Bioinformatics Platform, Monash University, Wellington Road, Clayton, Victoria, Australia; Biomedical Manufacturing, CSIRO Manufacturing, Bag 10, Clayton South, Australia; Department of Medical Biology, Academic Medical Centre, Amsterdam, The Netherlands; Faculty of Information Technology, Monash University, Clayton, Victoria, Australia; National Heart and Lung Institute, Imperial College London, London, United Kingdom; Systems Biology Institute Australia, Clayton, Victoria, Australia; University of Melbourne Centre for Cancer Research, University of Melbourne, Melbourne, Victoria, Australia

**Keywords:** Cardiac model, spatial transcriptomics, cardiac systems, bioinformatics, systems biology, 3D organ, data visualization

## Abstract

Understanding spatial gene expression and regulation is key to uncovering developmental and physiological processes, during homeostasis and disease. Numerous techniques exist to gain gene expression and regulation information, but very few utilise intuitive true-to-life three-dimensional representations to analyze and visualize results. Here we combined spatial transcriptomics with 3D modelling to represent and interrogate, transcriptome-wide, three-dimensional gene expression and location in the mouse adult heart. Our study has unveiled specific subsets of genes that display complex spatial expression in organ sub-compartments. Also, we created a web-based user interface for spatial transcriptome analysis and visualization. The application may be accessed from http://3d-cardiomics.erc.monash.edu/.

## Introduction

Organs or complex systems display a precise cellular spatial organisation which, if disrupted, can lead to functional changes and, eventually, disease. The mammalian heart is a complex organ composed of four structurally and functionally distinct chambers: left and right ventricles and atria (Moorman & Christoffels 2003), connected to the circulatory system through major vessels. It is composed of different cellular layers including cells types such as cardiomyocytes, fibroblasts, endothelial and immune cells (Massaia et al. 2018). This complex architecture ensures a plethora of functions such as contraction, electrical current conduction, blood and lymph circulation, and immune response. Different regions and structures of the heart exert different functions, hence molecular and physiological properties vary within the heart. For instance, intracardiac pressure is the highest within the left ventricle, which has the thickest muscular wall in the heart, to ensure blood distribution in the body. In contrast, the right atrium displays thin wall chambers as it only pumps blood to the lungs. Any anatomical or physiological alterations to these sub-compartments will impair cardiac function. For example, Hypoplastic Left Heart (Siffel et al. 2015), characterised by an atrophic or absent left ventricle, or transposition of the Great Arteries (Garne et al. 2007), manifested by an inversion of the connection of the pulmonary artery and aorta to the heart, are severe forms of cardiac malformations that require invasive surgery in the first years of life and can lead to death.

Our understanding of which genes are responsible for the formation and maintenance of specific cardiac sub-compartments is still limited. Thus knowledge of structure-specific gene expression is key if we want to address this. The importance of spatio-temporal gene expression and regulation in the heart has been well appreciated for decades (Waardenberg et al. 2014). Techniques such as is *in situ* hybridization or immunohistochemistry have shed light on the function of several genes in specific structures of the heart. However, this can only be done in a biased way and one or two genes at a time. In the last few years, different technologies to resolve spatial genome-wide expression in a systematic manner in whole organs or organism have emerged. For instance, spatial transcriptomics allows to interrogate gene wide expression in histological sections (Ståhl et al. 2016) (Asp et al. 2017) (X. Wang et al. 2018) as well as Slide-seq (Rodriques et al. 2019) and spatially barcoded arrays (Asp et al. 2017). DVEX (Karaiskos et al. 2017), Tomo-seq (Junker et al. 2014) and Geo-seq (Chen et al. 2017) have also been developed to capture different degrees of spatial resolution of gene expression in three dimensions. Recently, Burkhard and Bakkers utilised Tomo-seq to map the spatial transcriptome of the embryonic heart (Burkhard & Bakkers 2018). Although these techniques have enhanced our ability to determine and explore the spatial transcriptome of some model organisms and tissues, they have mostly been restricted to the size of the studied tissue, organ or organism. Consequently, to date, none of these methods have systematically investigated gene expression in 3D in adult mammalian hearts.

Here we present 3D-cardiomics, a three-dimensional gene expression atlas of the murine adult heart generated from RNA-sequencing of 18 anatomical sections. Analysis of this dataset revealed regional synexpression groups, including known cardiac-markers and novel compartment-specific genes. In addition, we present a novel visualization interface that facilitates interactive gene expression navigation, synexpression analysis and differential gene expression across sections. Our study provides a unique framework to explore gene expression in an adult mouse heart where information is scarce, and enables the identification of spatially-restricted genes at an unprecedented resolution.

## Results

### Revealing the spatial transcriptional profile of the murine adult heart

We aimed to evaluate the spatial transcriptional profile of the adult mouse heart. To achieve this, mouse hearts were isolated and microdissected in 18 anatomical sections. Microdissection of the heart consisted first in splitting the major vessels, atria and ventricles (Figure 1A). Three equally spaced transverse dissections were then performed on the ventricle, followed by further longitudinal dissections of the ventricles. High-throughput whole transcriptome sequencing (RNA-seq) was performed in duplicates for each of the 18 sections. Replicates highly correlate (Figure S1A) following batch effect removal (Figures S1B,C). In parallel, a 3D *in silico* model of the mouse heart (de Boer et al. 2011), (Aanhaanen et al. 2010) was digitally partitioned (Figure 1B) mimicking the 18 sections of the microdissected hearts (Figure 1C). RNA-sequencing data from each anatomical section was then mapped to its respective 3D partition. The software package Unity [https://unity.com/] was chosen to build the 3D-cardiomics tool due to its capabilities in operating 3D models. RNA expression values of the 18 pieces for each gene were mapped as colours on to the 18 virtual heart pieces previously dissected in silico. This allowed us to generate a digital transcriptome map of the composing sections of the adult mouse heart, explorable in three-dimensions, which we used to investigate spatial transcriptional networks within the heart.

**Figure 1:**
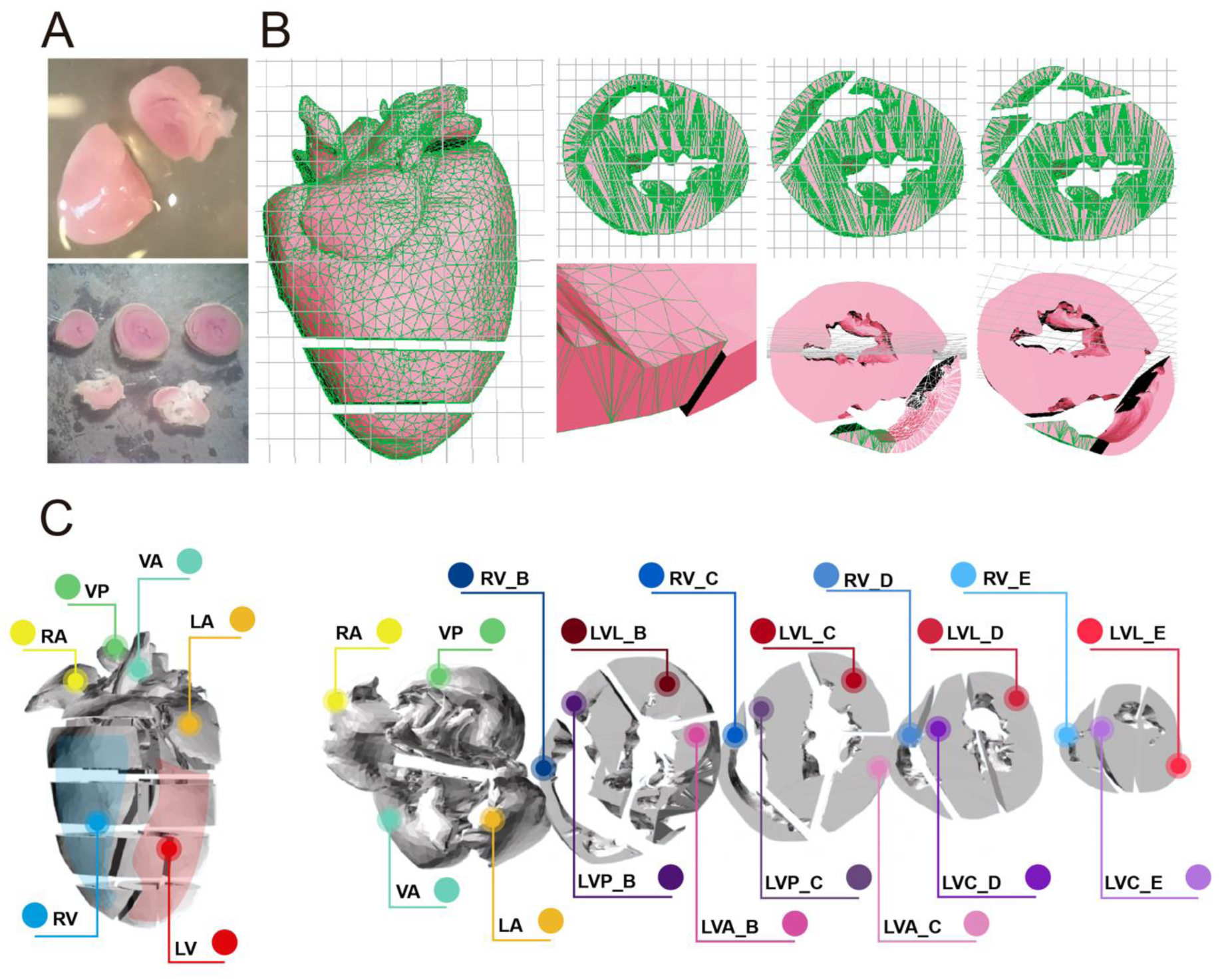
Schematic for designing the 3D-cardiomics interface, and nomenclature. A) Microdissection of the mouse heart. The atria and vessels were first sectioned from the ventricle (top), followed by sectioning of the ventricular pieces (bottom). B) *in silico* sectioning of the 3D murine cardiac model in Maya. C) Nomenclature for combinations of the specific sections and for each section of the heart model. Nomenclature: VP = vessel posterior, VA = vessel anterior, RA = right atrium, LA = left atrium, RV = right ventricle, LV = left ventricle, LVP = left ventricle posterior, LVA = left ventricle anterior, LVC = left ventricle centre, LVL = left ventricle left. The ventricle was sectioned into 4 larger sections which were B, C, D, E in order from the superior part of the ventricle to the inferior.

### Unravelling the structural transcriptome of the adult mouse heart

To gain insight into the spatial distribution of gene expression in the adult mouse heart, we examined how distinct cardiac sections clustered in low dimensional space. Correspondence analysis (CoA) revealed that the atria and major vessels diverged from the ventricular sections in the first two components (Figure 2A). The top 500 most variable genes of the first two components of the CoA explained most of the variability found between the cardiac sections and were associated with higher expression in either the atria or major vessels (Figure 2A). Investigation of variance explained by all components of the CoA revealed that most of the variability (62%) was explained by the first component if all sections were included in the analysis (Figure S2A). In contrast, the first component contributes only to 26% of the variance when only ventricular sections were analyzed (Figure S2B). To further explore the differences between the superior and inferior sections of the heart, we performed differential gene expression (DGE) analysis across the 18 sections, which confirmed the clustering of atrial and great vessel sections separately from the ventricular sections (Figure 2B). This is in accordance with the divergence observed from the CoA (Figure 2A). In order to spatially visualize the molecular segregation highlighted by the CoA and the DGE analysis, representative genes of each quadrant (Figure 2A) and cluster (Figure 2B) were visualized on our 3D *in silico* model (Figure 2C). The visualizations confirmed that the spatial spread of genes correlated with the spatial spread of sections in the first two dimensions of the CoA. A gradient across the second dimension was present which had separated the vessels from the atria. For instance, unbiasedly identified, *Tat* and *Uts2b* were highly expressed in the atria; *Bmp3* and *Gata5* have variable expression between the atria; *Adipoq* and *Pon1* are highly expressed in the major vessels. A noticeable gradient throughout the first dimension of the CoA, showed that the ventricles segregated from the superior tissues of the heart. The genes characterising the ventricles include *Irx1* and *Myl3* (Figure 2C). Some of these genes have been previously characterised in the specified sub-compartments, whilst others are novel candidate markers (Motoki et al. 2009) (Yue et al. 2017) (Gu et al. 2012), (Tward et al. 2002), (Shih et al. 1998), (Patel et al. 2008), (Andersen et al. 2012), (Zhi et al. 2016). Altogether, these findings confirmed that major spatial gene expression differences in the heart correlate with chamber identity.

**Figure 2:**
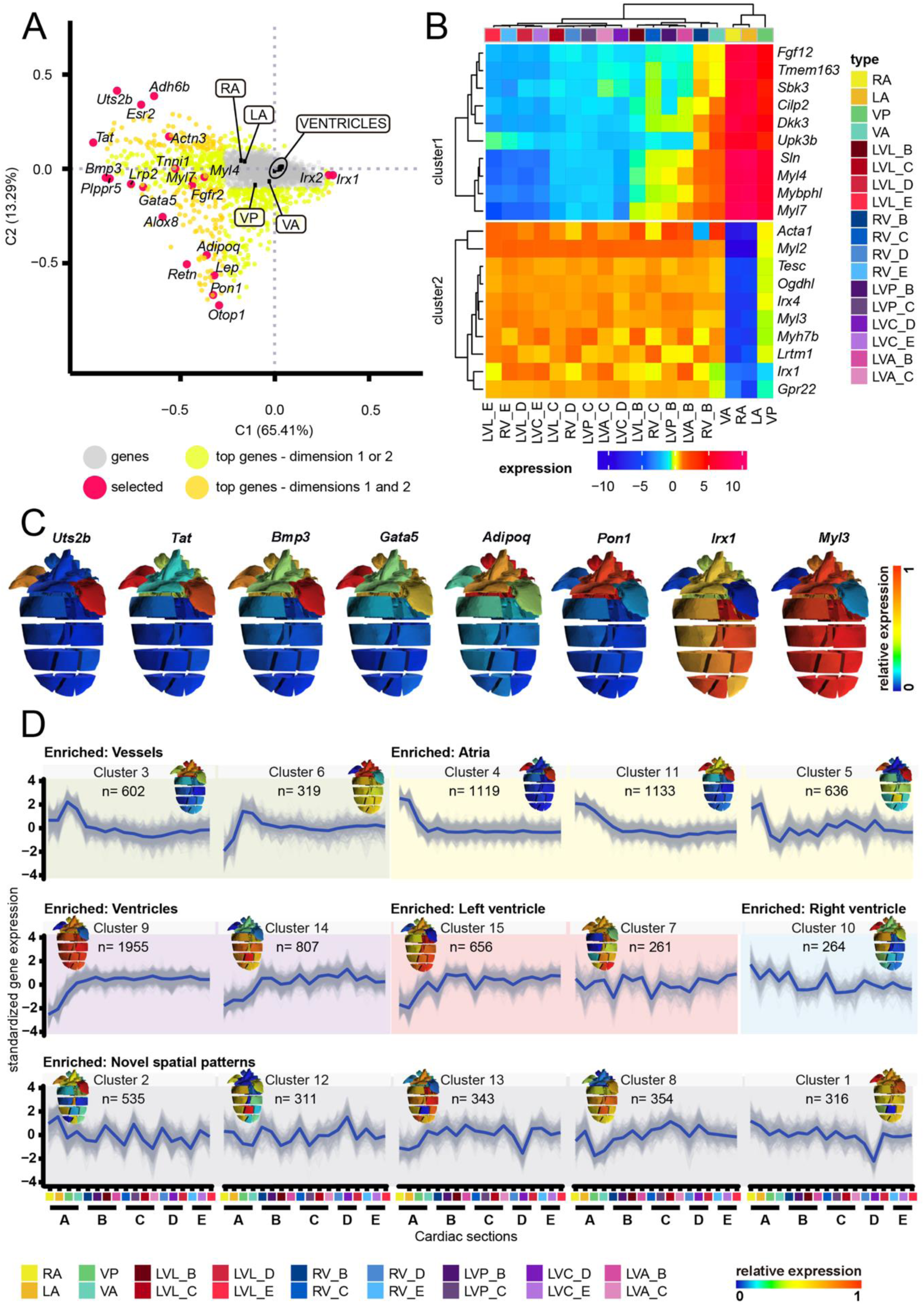
Characterization of gene expression across the mouse heart. A) Biplot, visualizing the separation of all genes and all sections in the first two components of CoA. Genes highlighted are the top 500 genes contributing to the most variance in components one and the top 500 genes in component two. B) Heatmap depicting hierarchical clustering of all cardiac sections and significant differentially expressed genes (atria and vessels against ventricles) with the highest absolute logFC values. C) Gene expression profiles of selected genes from the biplot visualized with the 3D-cardiomics user interface tool. D) Soft clusters of cardiac gene expression in mouse, represented in 2D and 3D.

To uncover global synexpression groups beyond these representative gene expression patterns, we performed unbiased soft clustering and 3D visualization across the heart (Figure 2D). The analysis revealed 15 clusters of which 10 had specific enrichment of gene expression localised to one anatomical compartment (*e.g.:* expression pattern restricted *to* vessels; atria; left or right ventricles). Interestingly, 5 clusters contained gene sets with complex expression patterns across anatomical sections (*e.g.*: expression pattern observed in atria, vessels and ventricular septum; atria and right ventricle) (Figure 2D). These novel synexpression groups indicate complex molecular functions shared across anatomical compartments (Table S2). Genes that are not spatially restricted (*i.e.*: highest probability of belonging to a cluster is less than 0.7) were clustered together (Figure S2C) and the observed average standardised expression was across all sections. Gene ontology (GO) analysis of these gene sets identified enrichment of cellular maintenance processes, supporting the role of this broadly expressed synexpression group in basic cellular functions (Figure S2D).

In summary, expected patterns of gene regulation in the murine heart were captured in our study and novel patterns were also revealed.

### Deciphering gene expression profiles of the atria

We next investigated the spatio-transcriptional changes amongst the atrial sections. Three of the previously identified 15 clusters revealed subsets of genes which were either up- (cluster 11, cluster 4) or down-regulated (cluster 9) in the atria relative to the rest of the cardiac sections (Figure 3A). Genes belonging to cluster 11 were found to be enriched in biological processes relating to extracellular structure, and those of cluster 4 were enriched in biological functions relating to cardiovascular development and GTPase mediated signal transduction. Unsurprisingly genes of the cluster associated to high gene expression in the ventricles was enriched in metabolic processes and mitochondrial function (Figure 3B). The distinct atrial specific expression is represented by known atrial markers (*e.g.*: *Myl4, Myl7 (Orr et al. 2016)*, (Huang et al. 2003)) and novel ones (*e.g.: Eps8*, *Usp11*), whereas *Myl2* and *Adra1a* are respectively known and unknown markers showing ventricular specific expression patterns (Figure 3C). We further validated *Myl4* in silico expression gene patterns by RNAScope *in situ* hybridization (Figure 3D). As expected, *Myl4* displayed high expression in both atria. We have also shown the DAPI negative control (Figure S2E), as well as an RNAScope *in situ* hybridization of *Ubc*, a highly expressed cardiac marker discovered in our analysis (Figure 3E, Figure 2SF) to confirm our findings. We also validated the expression of the cluster 9 gene *Adra1a,* and confirmed the transcript was restricted to ventricles (Figure S2H-I). Even though the relative expression of *Adra1a* was low, higher expression was still observed in ventricle, which demonstrated the sensitivity of our analysis.

**Figure 3:**
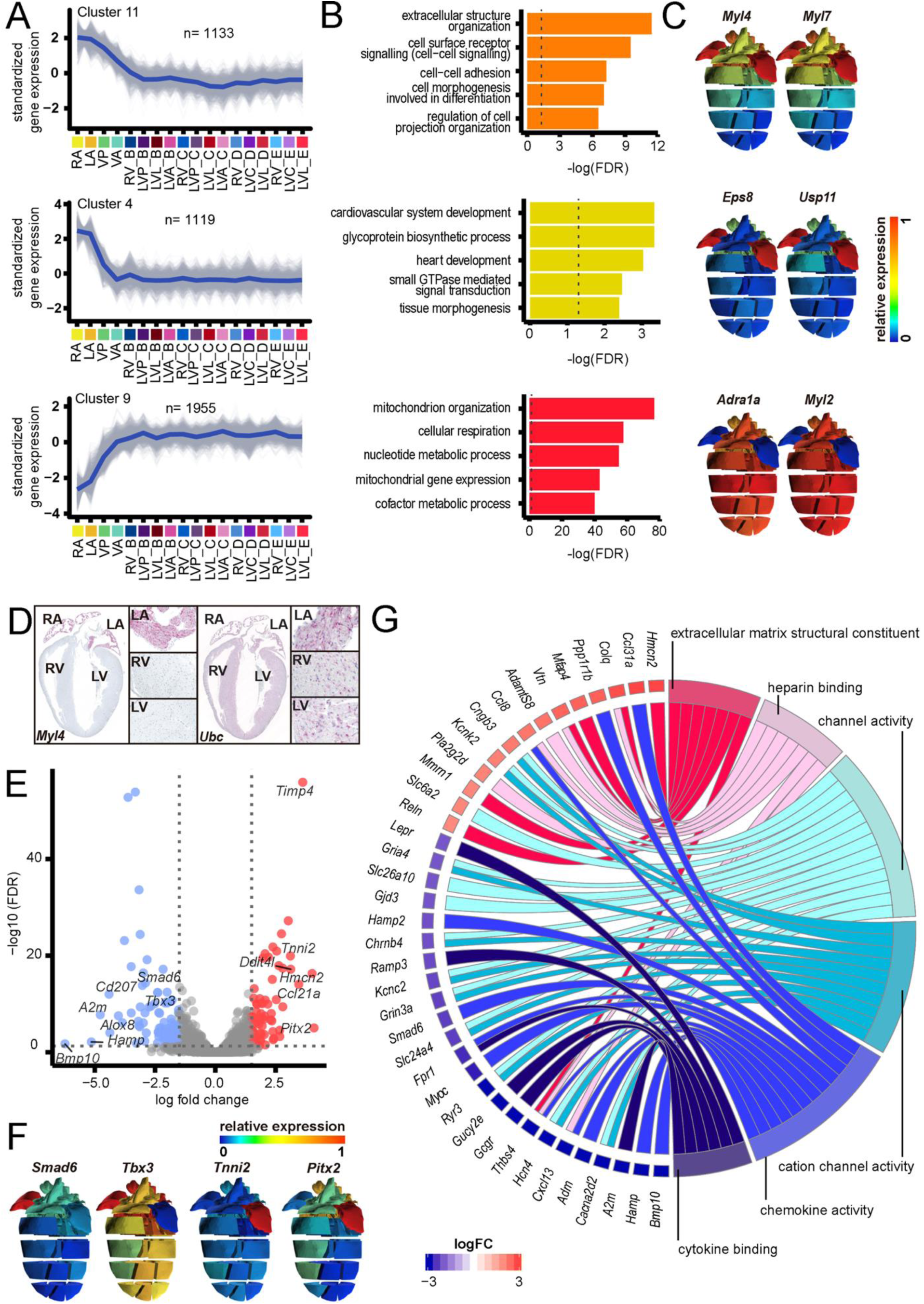
Transcriptome profiles of the atria. A) Soft clusters that contain genes describing the major differences between the atria and ventricles. B) The respective biological processes and C) selected gene expression profiles for soft clusters 11, 4 and 9. D) RNA-scope in situ hybridization of atrial specific gene *Myl4* which had been identified in clusters 4 and the highly expressed gene *Ubc*. Whole heart section is at 2.5X magnification and partial sections were at 10X magnification. E) Volcano plot of differentially expressed genes from DGE analysis of the left atria against the right. F) Gene expression profiles for several selected differentially expressed genes between the left and right atria. G) Circos diagram depicting differentially expressed genes with their log fold change, linked to one or more of the most enriched molecular functions of the DGE analysis.

We then investigated gene expression differences between the left and the right atria. No specific synexpression groups from the soft-clustering could capture differences between the atria. Thus, we performed a supervised DGE analysis between the left and right atria, and identified markers for each atrium (Figure 3E). The left atrium was characterised by known markers such as left cardiac lineage marker *Pitx2* (Campione et al. 2001) and the right atrium by the sinoatrial node transcription factor *Tbx3 (Hoogaars et al. 2007)* (Figure 3F). Consistent with this, KEGG pathway analysis of the identified DGE revealed enrichment of TGF-beta signalling, neuroactive ligand-receptor interaction and calcium signalling pathways in the right atrium (Figure S2F). In addition, comparative analysis of the molecular functions differentially recruited between the left or the right atrium suggested that extracellular matrix (ECM) functions (driven by well-known ECM components such as *Hmcn2*, *Adamts8*) are highly enriched in the left atrium (Figure 3G). In contrast, cytokine binding and channel activity was enriched in the right atrium (exemplified by markers such as *Lepr*, *Kcnc2*, *Hcn4*). Collectively, our findings provide insight into the specific transcriptional attributes of the atria where major expression differences pertain to the pacemaker functions restricted to the right atrium.

### Transcriptional complexity within the ventricles

In order to examine gene expression differences within the ventricular regions, we performed CoA on the ventricular sections exclusively (Figure 4A). Distribution of the section within the first two components revealed a tight clustering of the left ventricular sections away from the right ventricular sections, with the exception of the right, apical ventricular section (RV_E), which clustered closer to the LV sections. Soft clustering confirmed this distribution and identified two clusters containing genes which were highly expressed in the left ventricle (Figures 4B,C; clusters 15 and 7) and one cluster (Cluster 10) containing genes with higher expression in the right ventricle (Figure 4D). The CoA analysis also allowed us to determine which genes caused most of the variability in the ventricles (Figure 4A). These include *Nppb Efr3b*, *Brca1*, *Nrn1*, *Ces2e* and *Plekhh1* as enriched in the left ventricle and *Itga2b*, *Ngp* and *Tubb1* in the right ventricle (Figure 4E). Furthermore, as validation, we performed RNAscope *in situ* hybridization of *Nppb* expression (Figure S2K). Supervised DGE analysis between the left and right ventricular segments confirmed the findings using clustering analysis (Figure 4F). Indeed, we found a similar pattern of separation with the cardiac sections when we conducted hierarchical clustering using the genes with the highest log fold changes from our DGE analysis (Figure 4F). The genes with the highest log fold changes are common to the top genes from our CoA and soft cluster analysis, such as, *Itga2b*, *Tubb1*, *Nppb*, and *Plekhh1*. Intriguingly, we found that the top genes of the right ventricle regulate processes such as wound healing and blood coagulation (Table S1).

**Figure 4:**
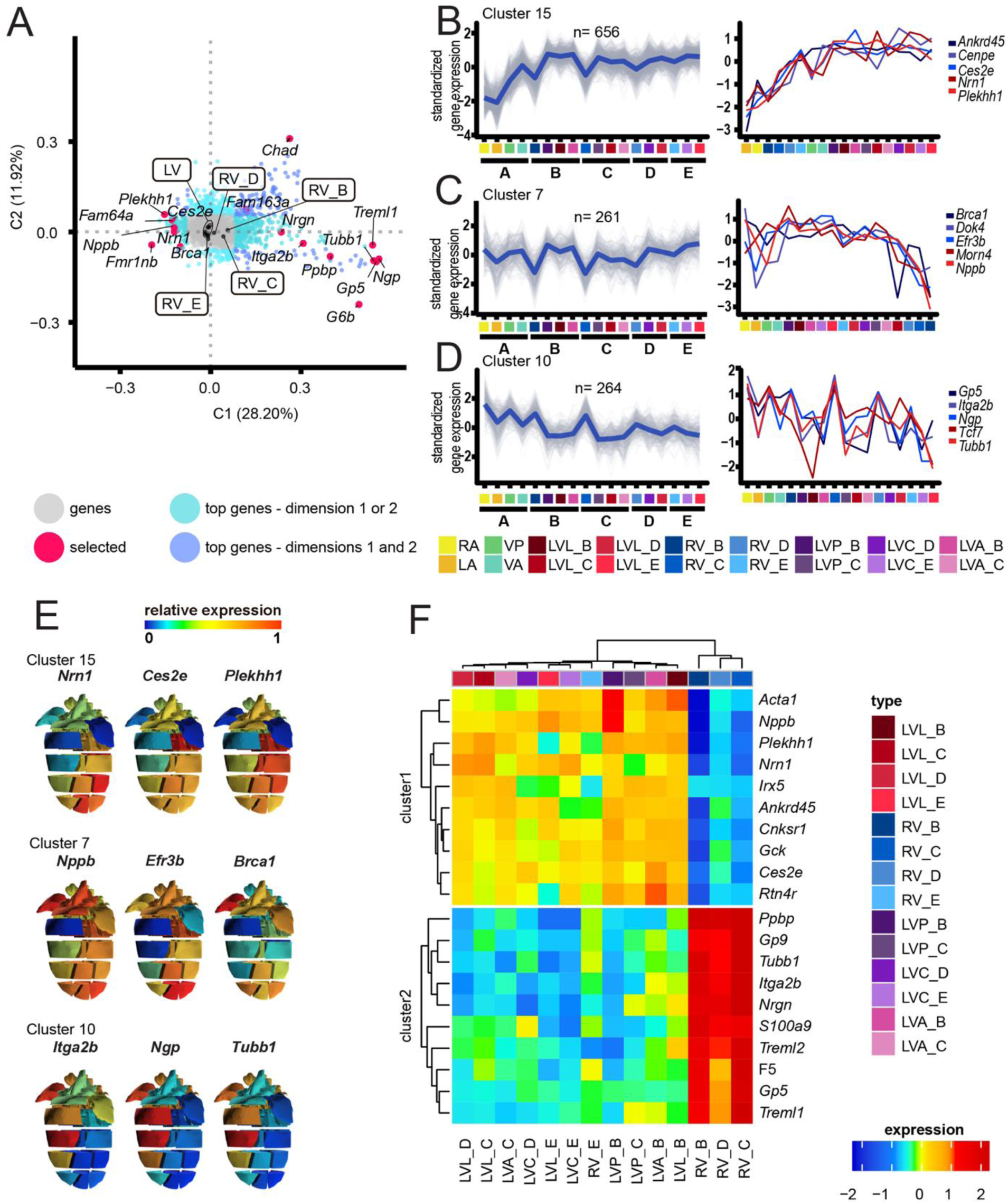
Transcriptional differences between the left and right ventricles. A) Biplot, visualizing the separation of all genes and all ventricular sections in the first two components of CoA. Genes highlighted are the top 500 genes contributing to the most variance in components one and the top 500 genes in component two. B-D) Soft clusters consisting genes highly expressed in the left ventricle (B,C) or the right ventricle (D), as well as gene expression profiles for selected genes from their respective cluster (sections of the selected genes plots were rearranged in the x-axis to clearly show the major differences between the ventricles). E) Selected gene expression profiles in 3D from respective clusters in B-D). F) Heatmap depicting hierarchical clustering of all cardiac sections and significant differentially expressed genes (right against left ventricle) with the highest absolute logFC values.

To further delve into the transcriptional complexity of the ventricles, we investigated additional sources of variation within the dataset. For this, we examined components 5 and 6 of the ventricular sections in the CoA (Figure 5A). We identified three clustered groups of sections corresponding to a unique spatial transcriptional pattern. These revealed that the inferior sections of the right ventricle (RV_C, RV_D, RV_E) segregate with the superior sections of the left ventricle (LVA_B, LVP_B, LVL_B). On the other hand, the superior section of the right ventricle (RV_B) clusters together with the remaining ventricular sections. The inferior section of the septum (LVC_D) surprisingly segregates away from both groups (Figure 5A). We then performed c-means clustering to further study these differences, which also revealed two similar clusters of complex patterns of gene expression across the two ventricles (Figures 5B). Interestingly both clusters show the inferior portion of the septum (LVC-D) to have very low gene expression in contrast to its neighbouring sections. This analysis provides a novel appreciation of the molecular differences within the two ventricles.

**Figure 5:**
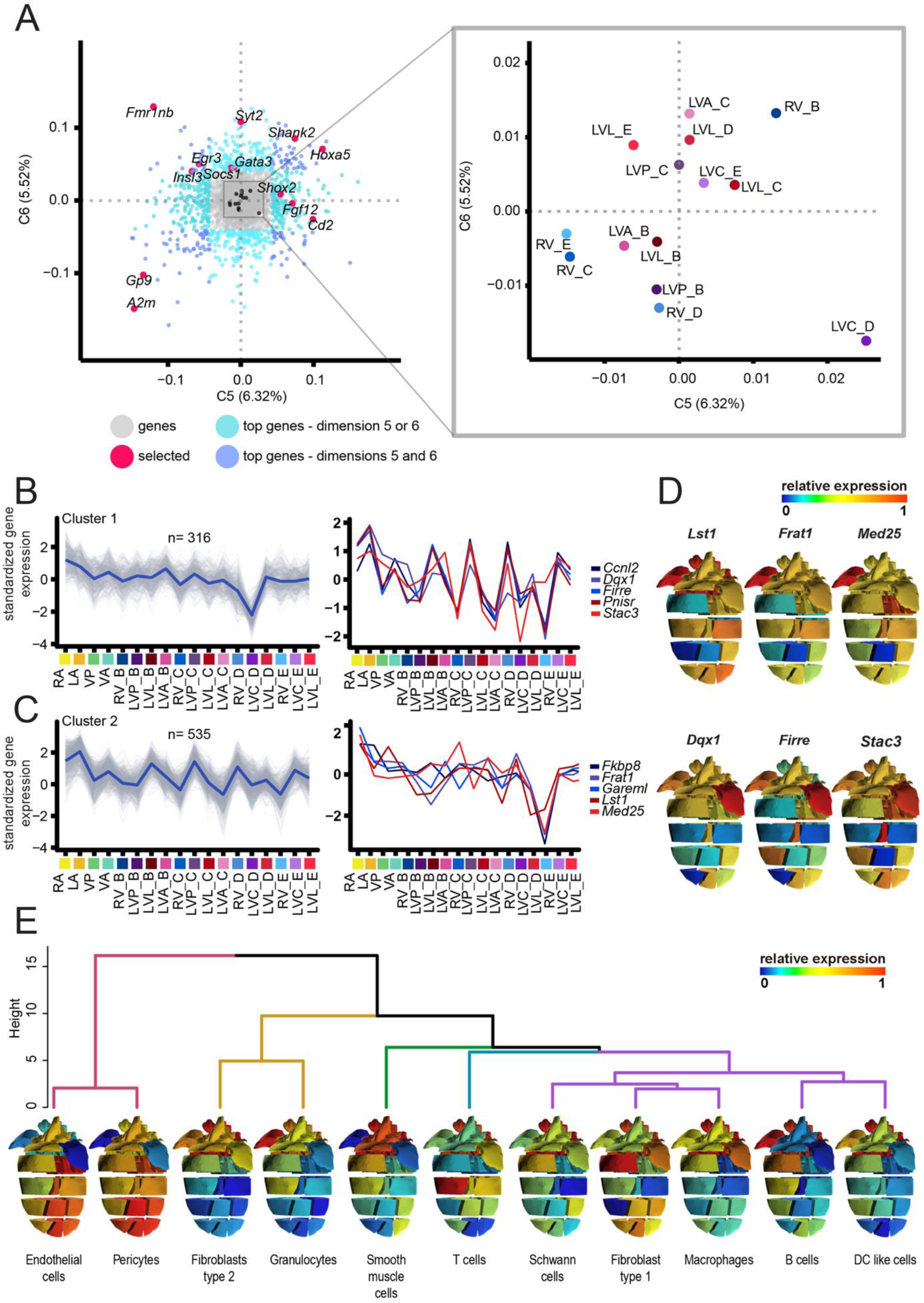
Further characterization of gene expression networks of the ventricles. A) The biplot which shows the spread of the ventricular sections, and genes in components 5 and 6. The biplot is enlarged at the coordinates of the ventricular sections to depict their spread. B-C) Clusters identified which depict an atypical gene expression profile of the cardiac sections, and individual gene expression profiles of selected genes within the identified clusters, which are also top genes in A). D) Selected gene expression profiles in 3D respective to clusters B and C. E) Hierarchical clustering of gene expression profiles of cell types across the heart.

### Prediction of non-myocyte localization within the heart

The novel spatial patterns discovered in this study lead us to investigate whether the molecular differences between the organ sub-compartments are due to their different functions and cell compositions. In order to address this, we mined the cell-specific markers unbiasedly identified by (Skelly et al. 2018) using their single cell analysis of the ventricles. We used the mean expression of markers from each cell type across all ventricular sections to generate a cell signature that we then superimposed in our 3D model. This in turn allowed us to locate where each cell type was most likely to be present in the ventricular sections and revealed that cell types are uniformly distributed across sections (Figure S2H). To further appreciate the spatial restrictions of different cardiac cellular subtypes in our 3D digital map, we repeated this procedure for the whole heart, performed hierarchical clustering and visualized the predicted locations of cell types within the heart (Figure 5E). Our analysis confirmed spatial enrichment of distinct cell-types in specific areas of the heart. For example, the spatial profiles of the two fibroblast subtypes differ from each other. Markers of fibroblasts type 2 display restricted expression in the vessels and atria, whereas fibroblasts type 1 are more widely distributed in heart. The most similar spatial expression profiles between two cell types are those of pericytes and endothelial cells, and of fibroblast 1 and macrophages. The novel spatial patterns of transcription identified, provide an insight into potentially new structural understanding of the organ.

### 3D-Cardiomics online tool allows for visualization of gene expression and differential gene expression analysis

In order to make the transcriptome analysis and visualization of the 3D model accessible to a wider audience, we created an online user interface which allows for further exploration of the data (Figure 6). The online tool includes features such as the interactive 3D heart, gene search and visualization, clusters from this study, and DGE analysis features (Figure 6A). In addition, a custom set of genes may be uploaded (for example single-cell markers), and the system will in turn extract a gene expression signature by performing an averaged expression of those genes for each section across the entire heart. The 3D model can be rotated or expanded for the examination of spatial patterns of gene expression (Figure 6B). The right panel displays this information for the current gene, sorted by the absolute descending value of the Pearson coefficient of determination (expressed as a percentage), giving the user genes which have RNA expression levels most correlated and inversely correlated with their current gene of interest (Figure 6C), allowing the identification of synexpression groups.

**Figure 6:**
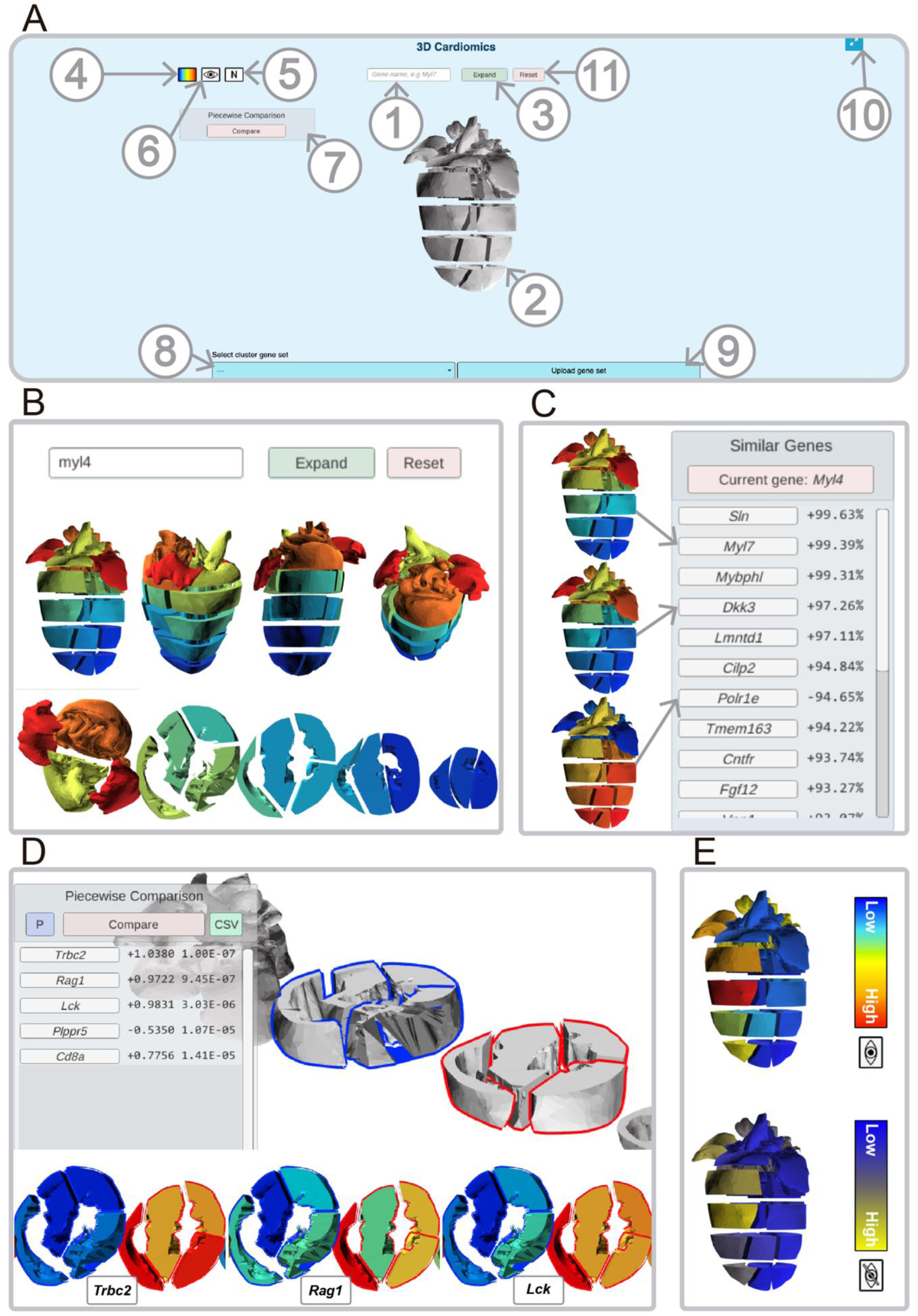
The interactive user web interface. A) Guide to components of the interface; 1) Search tool for genes expressed in the heart, for visualization, 2) 3D heart model, which displays gene expression profile of selected genes, 3) expand/collapse button for the 3D heart. 4) Colour based scale of gene expression, 5) Button which enables relative expression, providing a gene score ranging from 0 to 1, 6) color-blind color change option, 7) Cardiac section selector tool for pairwise differential gene expression, 8) Mean cluster (of this study) gene expression profile selection tool, 9) upload gene lists to obtain average signal across the heart, 10) enable full screen, and 11) reset analysis to collapsed heart, clear of gene expression. B) Visualisation of a selected gene. Top row shows the gene entered into the search bar, middle row shows different angles of the 3D model with the expression pattern of *Myl4,* bottom row shows the expanded view of the 3D model. C) Comparison of genes with similar expression patterns. Once a gene is selected a window appears with a list of genes highly correlated with the gene that was visualized at the time, with their correlation percentages listed. D) Example of differential gene expression analysis between ventricular segments A and B. A list of DEGs would appear with their LFC values and FDR values. ‘CSV’ is an option of downloading the output. When genes are pressed the expression patterns can be visualized such as the examples shown. E) Visualization of expression patterns in the different colour formats. Top shows the default, and bottom shows the color-blind friendly option.

For the first time, a 3D interface for exploring spatial gene expression offers real time differential gene expression analysis. Comparisons between any two sets of cardiac sections can be performed “on-the-go” and directly visualized by the “Piecewise Comparison” panel of 3D-cardiomics. Selecting “Compare” allows the input of the first set (which may include more than one sub-compartment), which can be selected by clicking on the model (Figure 6D). Finally, data can be visualized using two colour modes (Figure 6E) facilitating the website accessibility. In summary, our tool offers an ease of analysis and visualization of the adult mouse transcriptome, which could be applicable to any other organ or tissue 3D model.

## Discussion

Our study presents a novel way to integrate high-throughput data on a three-dimensional model for analysis. We have demonstrated an alternative and beneficial method for data visualization of gene expression data in 3D of an organ, alongside conventional approaches. For example, in our soft cluster analysis, we have provided the standardised gene expression line graphs concurrent to the cluster expression means visualized on the 3D heart. Similarly we have also provided the localisation predictions of numerous cell types in both the 3D model and via a heatmap. In both of these cases, the contrasting expression profiles between the clusters, or the different cell types is evident in both visualization approaches, however, the 3D heart poses as a more intuitive form for visualization in the complex spatial context.

The online tool we have created is advantageous, as expression profiles of sufficiently expressed genes or groups of genes in the heart can be visualized, as well as corresponding correlated genes. Additionally, our tool effortlessly allows for differential gene expression analysis, and visualization. Furthermore, the easily accessible tool can be freely used for visualization and analysis of cardiac related gene expression, which is beneficial for hypothesis building and uncovering of new roles of gene expression in the mammalian heart.

Our spatial transcriptomic approach allowed us to show that most of the variability between the cardiac sections is due to the difference between the atria, major vessels and ventricles. In the major vessels, we found enrichment of genes functionally associated with metabolism. For instance, we found *Pon1,* a gene known to be associated with atheroprotective effects and regulation of cardiovascular disease (Tward et al. 2002), (Shih et al. 1998). Similarly, *Adipoq*, which is also known to affect the artery intima media thickness (Patel et al. 2008), displays high expression in the major vessels. In conjunction with our findings, *Adipoq* is also present in aortic endothelial cells (Komura et al. 2013), hence this three-dimensional model serves well for hypothesis testing of other potentially present proteins previously not associated with any of the cardiac sub-components. We also uncovered known and unknown markers for the atria and ventricles, such as *Myl4* or *Myl3* respectively (Gudbjartsson et al. 2017) (Peng et al. 2017) (Orr et al. 2016). *Myl3* has been known to be associated with all sub-compartments of the heart, however the degree of differential expression across compartments had been masked by the lack of spatial information (T. Y. Wang et al. 2018). Differential gene expression analysis between left and right atria revealed enrichment of *Tbx3* expression in the right atrium, a crucial transcription factor governing the establishment of the sinoatrial node, also located in the right atrium. Not surprising, the most distinct ventricular pattern was between the left and the right chambers. The left ventricle was found to be enriched in functions associated to muscle regulatory processes such as higher respiratory and muscle activity which is concordant to previous research, as the left ventricle is required to supply blood to the entirety of an organism. In contrast, the right ventricle pumps blood through, to the pulmonary vasculature, and only with about 25% of the stroke work of the left ventricle (Sordahl 1976) (Voelkel et al. 2006).

Beyond these major differences between atria and ventricles, we identified a unique set of spatial gene expression signatures. We hypothesised that the cellular composition between the sub-compartments contribute to the observed differences. To aid in our understating of the dominant cellular compositions retaining spatial information, we exploited our 3D model to map cell-type specific gene signatures across the heart. Interestingly we found endothelial cells and pericytes to have an approximately equivalent spatial distribution. The concurrence of these two cell-types has been extensively established (Hellström et al. 2001) (Franco et al. 2011) (Kato et al. 2018) (Murray et al. 2017), hence it is captivating to find their 3D profiles correlate within the heart. This expected and well supported finding is a positive control for the additional cell-specific signatures that we have revealed. Indeed, our single cell localisation hypotheses also suggested that T cells and B cells should be enriched in the midline-transverse section of the right ventricle. Studies have previously shown that the heart is comprised of cells from the hematopoietic lineage, however this novel link to a spatial compartment uncovers new possibilities for the roles of these cells in the heart (Farbehi et al. 2019), (Skelly et al. 2018), (Holzinger et al. 1996), (Pinto et al. 2016). This model holds as a suitable benchmark for hypothesis testing of cardiac cellular localisations, and possible functions of the various anatomical components.

Curiously, there were patterns indicating septal divergence from our CoA and cluster analyses, in particular the section LVC-D appeared to have dissimilar expression to its neighbouring sections. These findings align with some of the cell-type specific spatial profiles, which could account for the distinction of LVC-D, in particular Schwann cells, the subset of fibroblasts identified by (Skelly et al. 2018) fibroblasts 1 and Macrophages. This difference, for instance, could be in consequence of the discontinuity of the glial cell populations, which pass down through the septal region and then diverge in the central region to the rest of the heart as Purkinje fibres (Anderson et al. 2009).

We have also uncovered upregulated genes in the superior region of the heart in comparison to the other ventricular sections. It was riveting to discover that fibroblasts 1 and macrophages had a hypothesised concentration in the apex of the heart, as well as an almost identical spatial profile. In the context of cardiac regeneration this is very enticing as both fibroblasts and macrophages are known to communicate extensively post myocardial infarcts (MI)s and jointly play a substantial role in cardiac repair, by promoting debris clearance and the establishment of the fibronectin matrix. (Sattler & Rosenthal 2016) (Chen et al. 2012) (Forte et al. 2018). It would be very intriguing to find that their communication is required for homeostatic cardiac function based on the evidence from our study, however this is something yet to be investigated.

Having drawn these conclusions, it is necessary to state that there are limitations in single cell studies of the heart, which impacts our hypotheses on cardiac cellular localisation. (Skelly et al. 2018) have only identified two subtypes of fibroblasts, however, (Farbehi et al. 2019) have identified more subtypes in the adult mouse heart. The reason for this may be because more rare cell types were captured in (Farbehi et al. 2019)’s study as only the apical section of the heart was isolated, in comparison to (Skelly et al. 2018)’s, which utilised the entire ventricle. This shows that the use of the 3D-cardiomics tool, and hypotheses drawn from it could be subjective to prior analyses. Having the spatial information of cell types throughout an organ tells us not only more about the role that these cells may play in a homeostatic setting of the heart and how they maintain the organ, but we could also gain a better understanding which cells function or communicate together, and hence possibly further our understanding of these roles in other contexts such as cardiac injury and repair.

In summary, we propose that by retaining the spatial signature of the transcriptome of organs and in combination with 3D models not only allows us to visualise expression patterns across an organ but greatly enhances discovery. We anticipate that the capacity of 3D-Cardiomics to be combined with single cell or pathological signatures will be of great utility to the cardiac field.

## Materials and Methods

### Key resources table

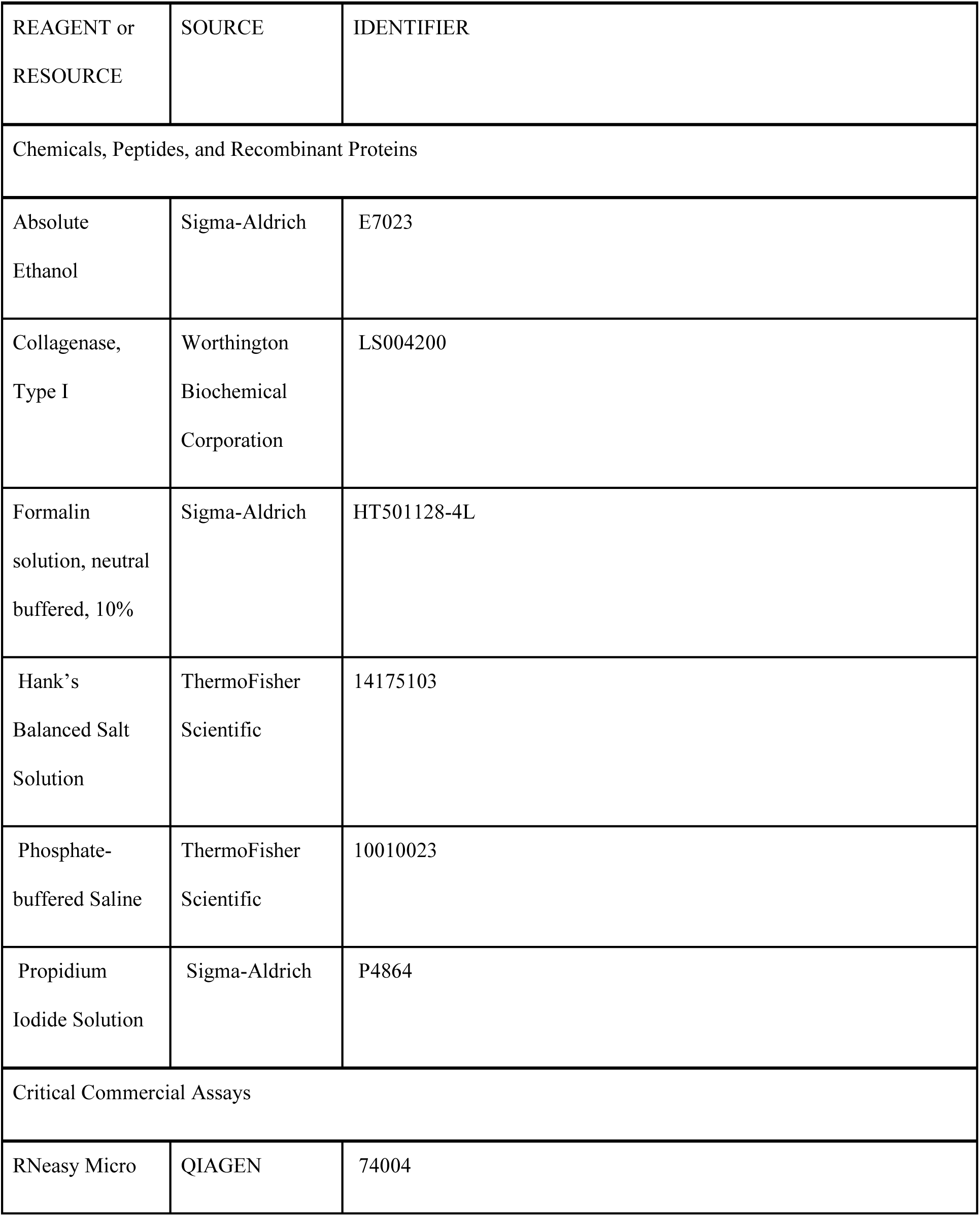
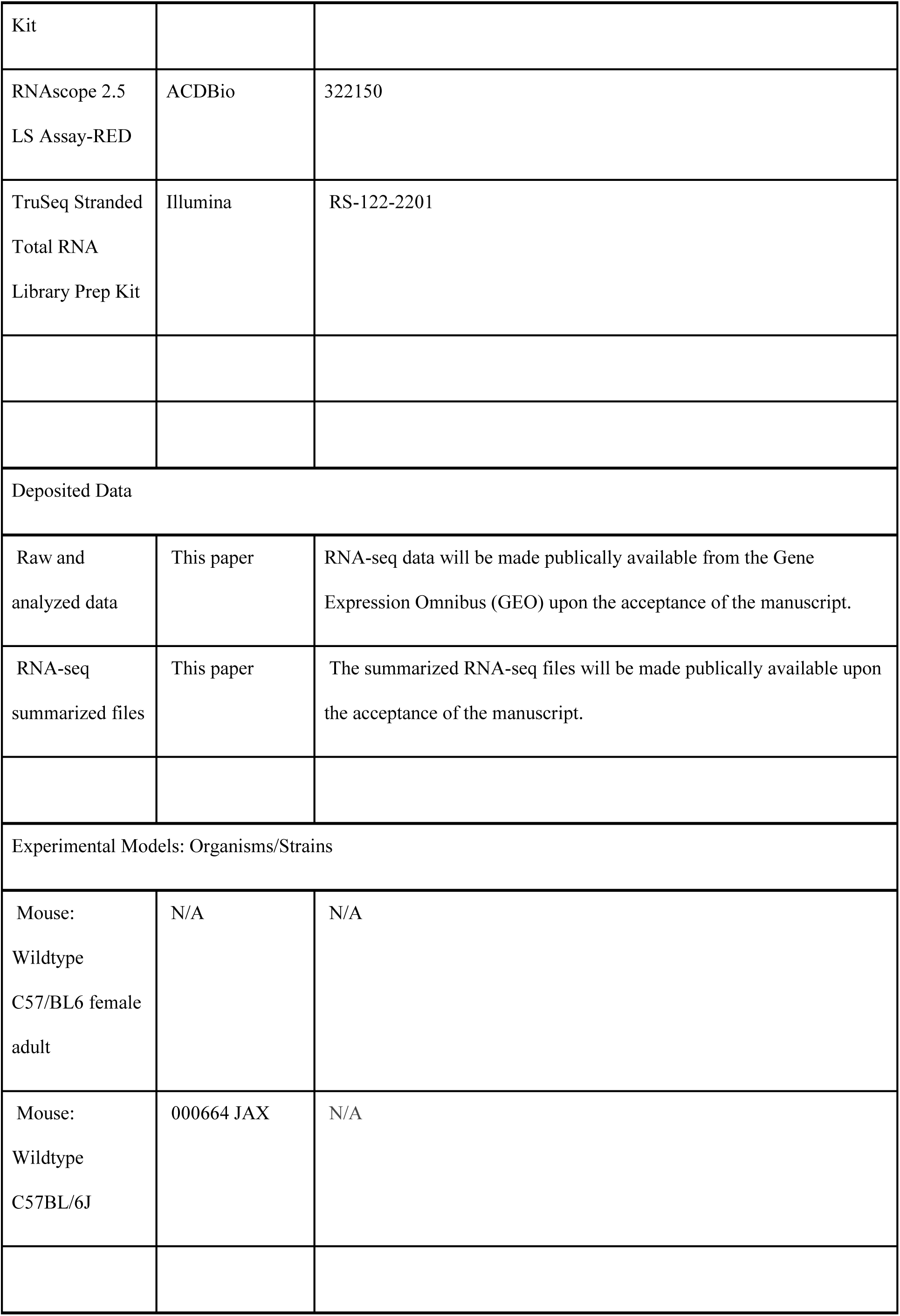
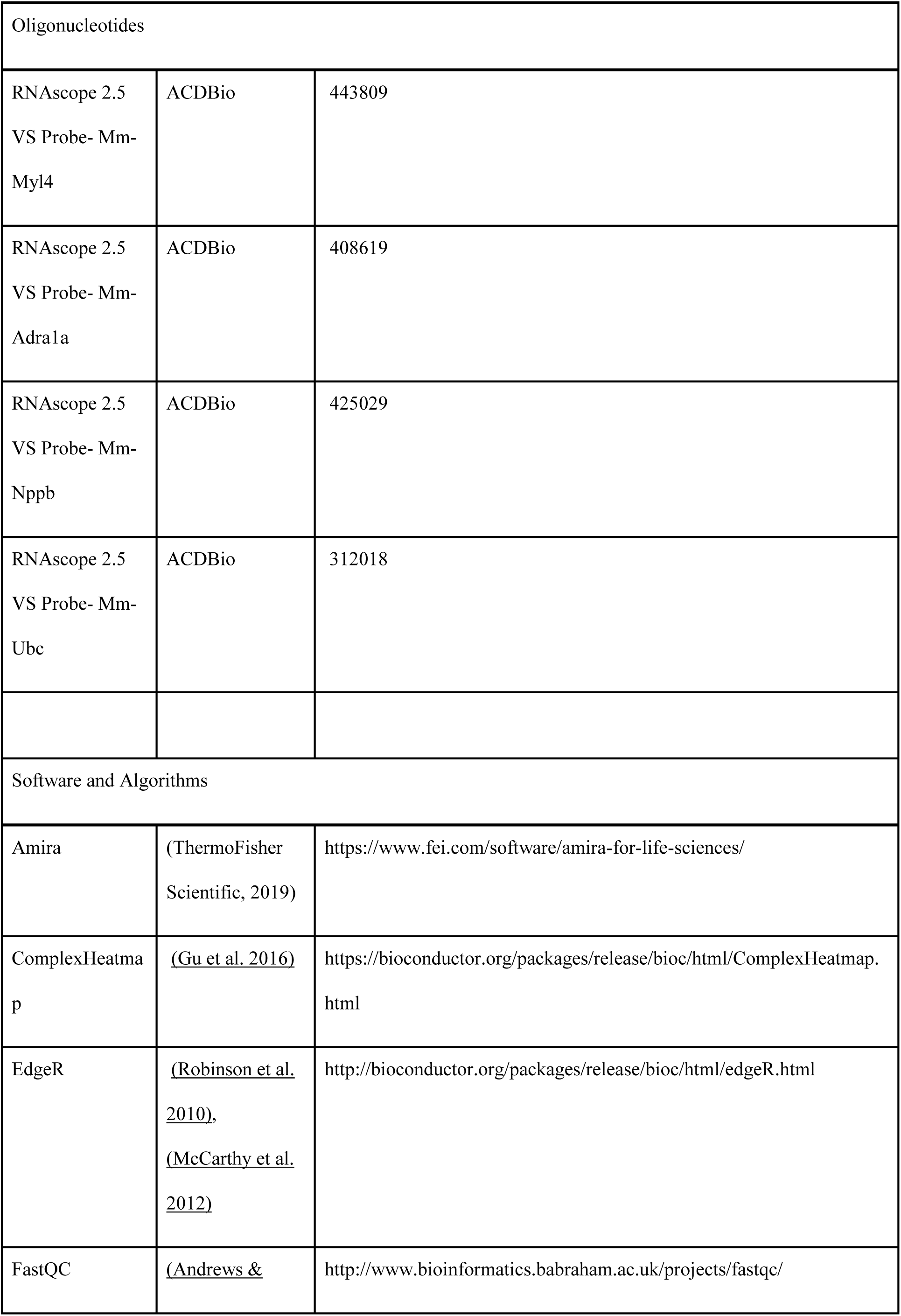
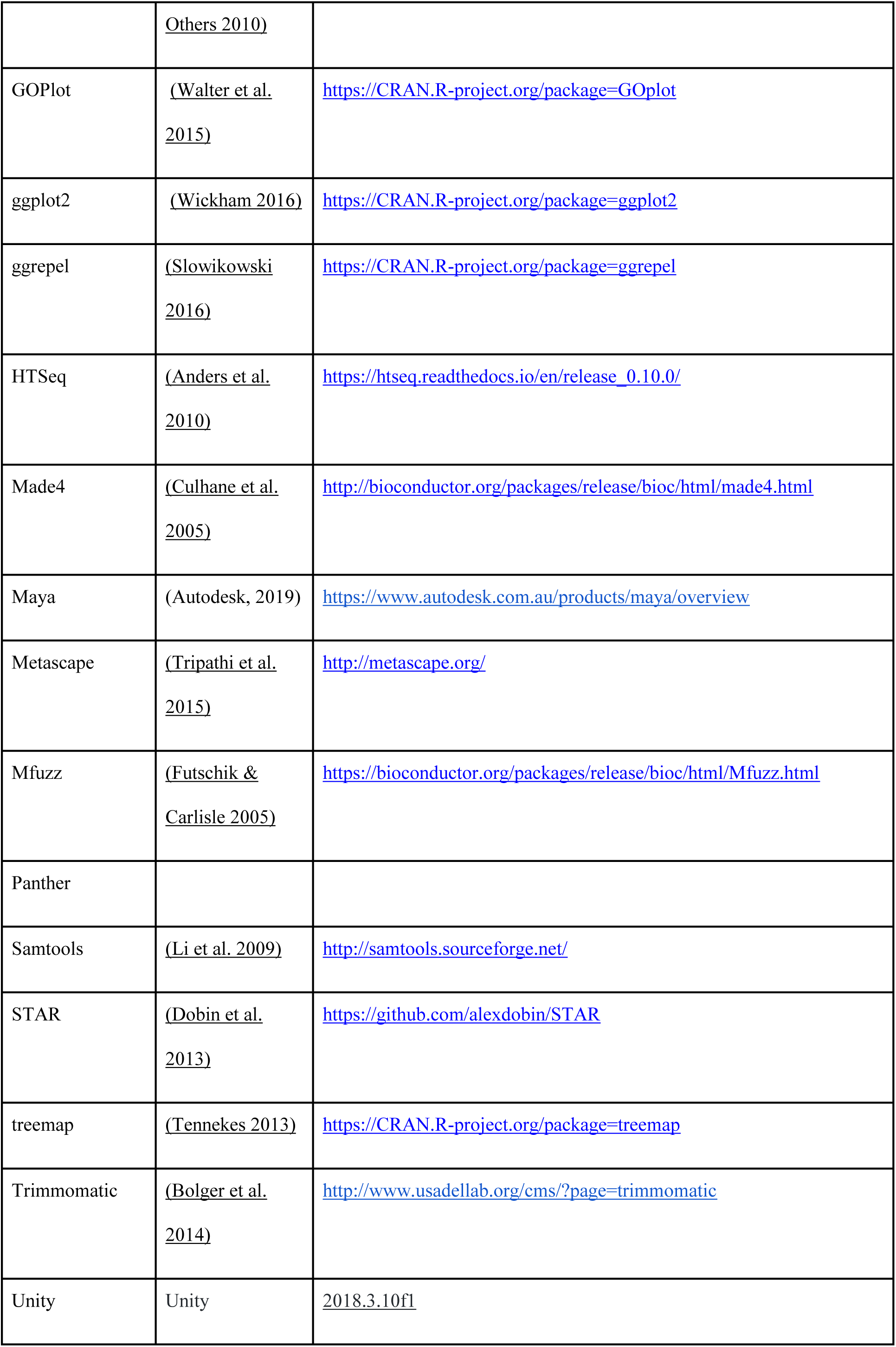
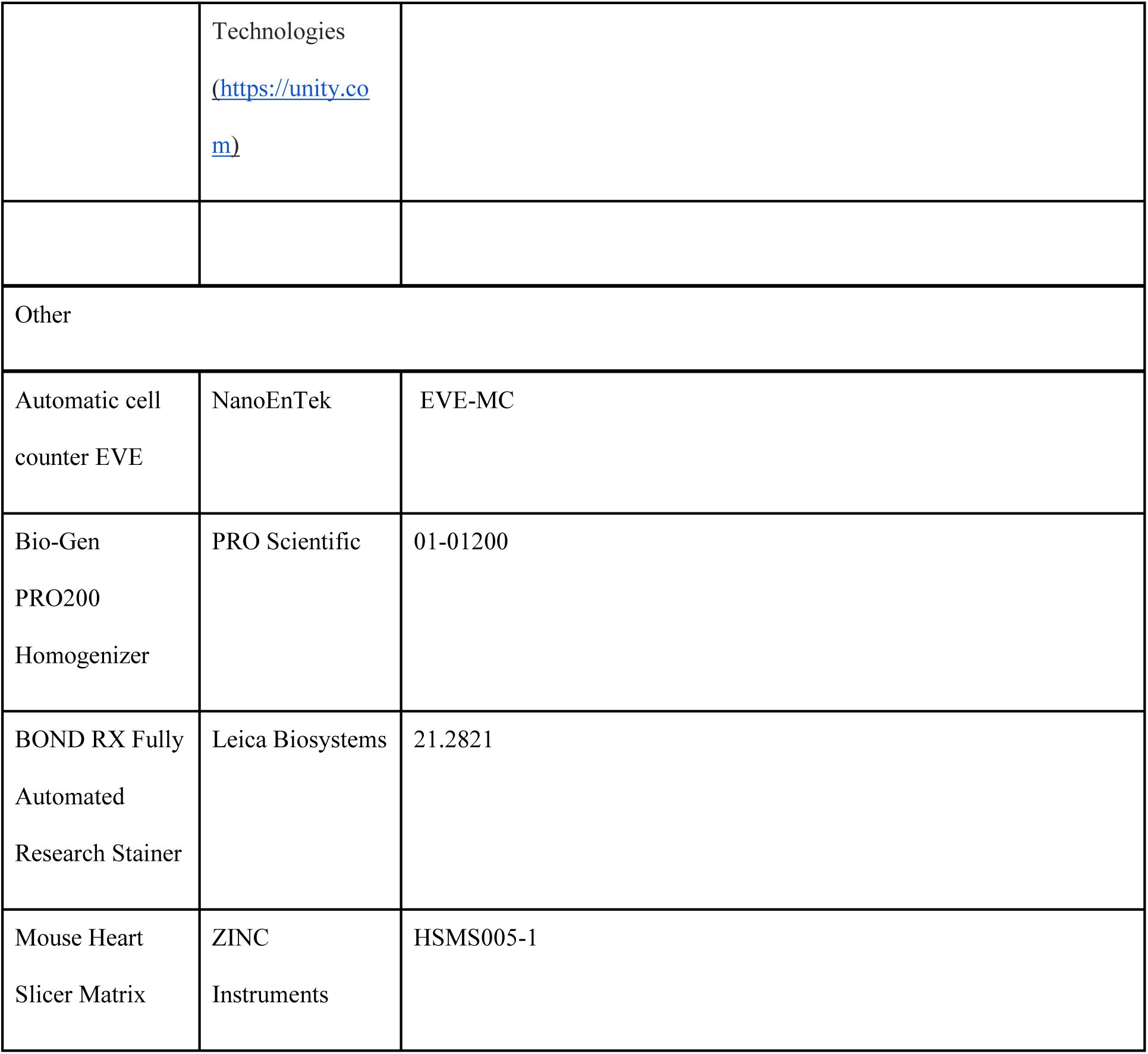

### Contact for reagent and resource sharing

Further information and requests for resources and reagents should be directed to and will be fulfilled by Lead Contact Jose Polo.

### Experimental model and subject details

#### Mice

For the sequencing experiment, C57/BL6 mice were housed at the Monash University animal facility in strict accordance with good animal practices defined by the National Health and Medical Research Council (Australia) Code of Practice for the Care and Use of Animals for Experimental Purposes. All experimental procedures were performed under the approval of the Monash University Animal Research Platform animal ethics committee. For the *in vivo* validations, C57BL/6J from JAX, were used with good animal practices defined by the Public Health Service Policy on the Humane Care and Use of Laboratory Animals and the Guide for the Care and Use of Laboratory animals; in compliance with all federal, state and local guidelines, regulations and animal care programs, fully accredited by the Association for the Assessment and Accreditation of Laboratory Animal Care International.

### Method details

#### Heart extraction and microdissection

Mice of approximately 6-10 weeks of age were culled using the cervical dislocation method, then sprayed with 80% v/v ethanol and immediately dissected through the abdomen under the sternum. Once the diaphragm was dissected away to access the upper abdominal cavity, the rib cage was cut and lifted to expose the heart and lungs. Perfusion was then performed on the heart. Firstly small incisions were made on each lobe of the liver to aid in bloodletting. Hanks’ Balanced Salt Solution was delivered through the left ventricle for one minute, using a 10 mL syringe with a 26-gauge needle. Hearts were removed by grasping the heart by the root and cutting through the major vessels and surrounding connective tissue. Once removed, the heart was placed in 310 mO Phosphate Buffered Solution (PBS) to clean up excess fat, connective tissue, lungs thymus and trachea.

The isolated hearts were first dissected using Vannas scissors to remove the atria and major vessels. Each ventricle was then microdissected into four equidistant transverse sections with a mouse heart slicer matrix. An anchor blade was used at the superior end of the heart to ensure the heart did not move. The blades were firmly pressed down simultaneously to cut through the ventricles. All sections were then placed into PBS and individually dissected into the sections as specified in Figure 1C with Vannas scissors.

#### Tissue preparation

The microdissected sections of each heart were placed in 250 𝞵l Buffer RLT as supplied by QIAGEN RNeasy Micro Kit. Six samples were processed at a time whilst the remaining sections were stored at −80°C. The samples were homogenised for 30 seconds at medium speed (setting 2). The homogeniser was cleaned with 80% v/v ethanol for 15 seconds, and then with deionised water for 15 seconds. The samples were then immediately processed for RNA extraction.

#### RNAScope in situ hybridization

Hearts were fixed in 10% neutral buffered formalin (Sigma-Aldrich), processed for paraffin embedding and 5µM longitudinal sections used for automated staining with RNAscope 2.5 LS Reagent Kit—Red (ACDBio) on a Leica Biosystems’ BOND RX Research Advanced Staining System (Leica). Probes used were as follows: Myl4 443809; Adra1a 408619; Nppb 425029 and positive control Ubc 312018.

#### RNA sequencing

RNA extraction and purification of all samples was done using the RNeasy Micro Kit according to manufacturer’s instructions. An illumina TruSeq Stranded Total RNA kit was used to prepare the libraries for poly-A enrichment of mRNA. For sample amplification, 15 PCR cycles were used, standard to the Illumina kit protocol. RNA sequencing was performed on an Illumina NextSeq500. Each library was paired-end with 75bp reads, as well as 20 million reads per sample.

#### 3D Heart Model

Amira was used to export the 3D graphical model of the adult mouse heart [reference Ruijter] into Wavefront .obj files, which could then subsequently be used with 3D modeling software Maya (2015) to perform the computational slicing and sectioning. For this, the heart was first sliced into 5 transverse pieces as with the biological samples. Slicing of the 3D model was done using the “Slice” tool in Maya, which allows a straight line to be drawn to cut the object completely through. Each slice was then ‘sealed’ to give the appearance of solid tissue. The ‘sealing’ of the slices was done by adding faces individually to the model until the slice was completely covered. Once sealed, each slice was then sectioned into 18 pieces, again using the Slice tool. Similar to the slicing process, each piece was subsequently sealed by adding faces.

#### User interface

A visual system was developed to integrate RNA-seq datasets onto computational model pieces of the heart using the C# programming language in the Unity environment. The program was compiled to the WebGL platform to allow cross-platform accessibility through a web browser and fast data retrieval. Source code is freely available on GitHub (https://github.com/Ramialison-Lab-ARMI/3DCardiomics). For visualization purposes, the normalized values of gene expression observed on the web interface were calculated by normalization to the local minimum and maximum expression of the gene.

#### Gene expression analysis

Raw sequencing reads were filter/trimmed using trimmomatic (Bolger et al. 2014). Sequencing reads were aligned to GENCODE’s mouse reference genome (GRCm38 primary assembly, vM9 annotation) with STAR (v2.4.2a), (Dobin et al. 2013). Gene read counts were generated with HTSeq (Anders et al. 2010). Genes with 1 count per million (CPM) in at least two of the samples were kept for further analysis. For differential expression analyses, normalisation factors were calculated by the trimmed means method using the EdgeR function calcNormfactors (Robinson et al. 2010). The glmFit and glmLRT functions from the EdgeR package were used to perform differential gene expression analysis. The design matrix required for these analyses included specification of heart segment and batch label, in order to remove the batch effect. For the remainder of the analysis CPM values were used throughout the study, and RPKM values were used for the cluster analysis. To remove the batch effect for further analyses removeBatchEffect from EdgeR was used. The plotMDS function from EdgeR was used for MDS analysis and visualisation. Made4 was utilised for CA (Culhane et al. 2005). Mfuzz was used for the soft cluster analysis. The number of clusters specified were determined by the total number of dimensions explaining at least 95% of the variance in the data from the COA on the dataset with RPKM values. Metascape was used for biological processes enrichment. The treemap package was used for the treemap visualizations. All visualizations were made with ggplot2 unless otherwise specified. Unsupervised hierarchical clustering and the ComplexHeatmap map package were used on significantly differentially expressed genes.

## Acknowledgements

We would like to acknowledge the Histology and Microscopy Cores of the Jackson Laboratory for performing the RNAScope experiments, the Ramaciotti Centre for Genomics for performing the NGS experiments. We would also like to thank the Monash Animal Research Platform and Monash Flowcore for providing required services throughout the course of the project. J.M.P. was funded by a Sylvia-Charles Viertel Fellowship and an ARC Future Fellowship M.R. was funded by a NHMRC/Heart Foundation Career Development Fellowship and ARC Discovery Project Grant. A.T. was supported by a Biomedical Research Victoria UROP/CSL scholarship. The Australian Regenerative Medicine Institute is supported by grants from the State Government of Victoria and the Australian Government. Finally we would like to thank Dr. Jan Ruijter, Dr. Alexander Pinto, Ms. Jeannette Hallab, Mr. Markus Tondl, Dr. Michael Eichenlaub, Dr. Anja Knaupp, Dr. Ethan Liu, Dr. Jaber Firas, Dr. Sue Mei Lim, Dr. Sara Alaei, Dr. Gonzalo Del Monte-NIeto, the Ramialison laboratory and the Polo laboratory for their valuable contributions to this study.

## Author Contributions

J.M.P. conceived the study. J.M.P. and M.R. designed the experiments and co-lead the project and supervised the project together with F.J.R.. M.M. and F.J.R. performed the bioinformatics analysis, with contributions from N.M.T., A.T. and H.N.. M.M. interpreted the data with contributions from N.T, A.T.. M.M. generated the figures with contributions from N.M.T., A.T., M.B.F., M.W.C. and F.J.R.. N.T. performed the heart isolation with input from J.H., S.K.N., M.F. and M.W.C. and generated the transcriptional data and adapted the K.V.D. 3D model of the heart. A.T. developed and optimised the 3D-cardiomics user interface with contributions from N.M.T., A.P., D.P. and F.J.R.. M.F., M.W.C. generated the RNAscope datasets. M.M., F.J.R., M.R. and J.M.P. wrote the manuscript with contributions from N.M.T., A.T.. All authors approved of and contributed to the final version of the manuscript.

## Supplemental Information

### Supplementary table legends

**Supplementary table 1: CA and DGE analyses outputs**

The top 500 genes explaining most of the variability in the data of the CA, which included all cardiac sections (sheet 1), with enriched biological processes (sheet 2); the top 500 genes of the CA analysis, which included ventricular sections, in dimensions 1 and 2 (sheet 3) and respective biological processes enriched (sheet 4); the top 500 genes of the CA analysis, which included ventricular sections, in dimensions 5 and 6 (sheet 5) and respective biological processes enriched (sheet 6). Finally, the DGE analysis outputs of the left atria compared to the right (sheet 7) and the right ventricle compared to the left (sheet 8).

**Supplementary table 2: Soft cluster analysis outputs**

Outputs of the soft cluster analysis. Firstly, a list of all of the genes belonging to a cluster with a probability of at least 0.7 (sheet 1). The remaining sheets include biological processes enriched for each of the clusters.

---

**Supplementary figure 1:**
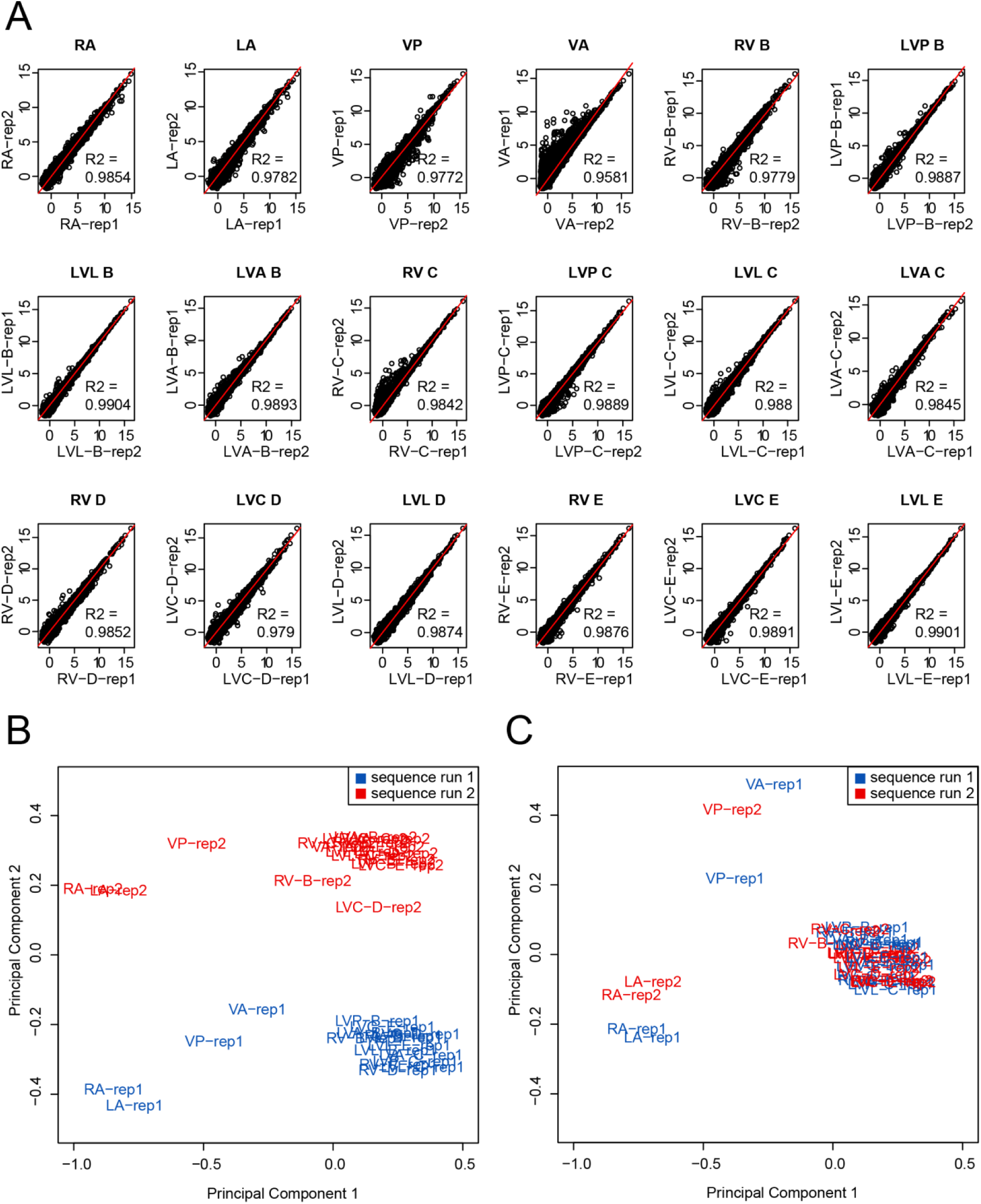
Replicate correlation measures and batch effect removal. A) The correlation plots of counts of each gene for each of the replicates. B) Principal component analysis of all samples before batch effect removal and C) after batch effect removal.

**Supplementary figure 2.**
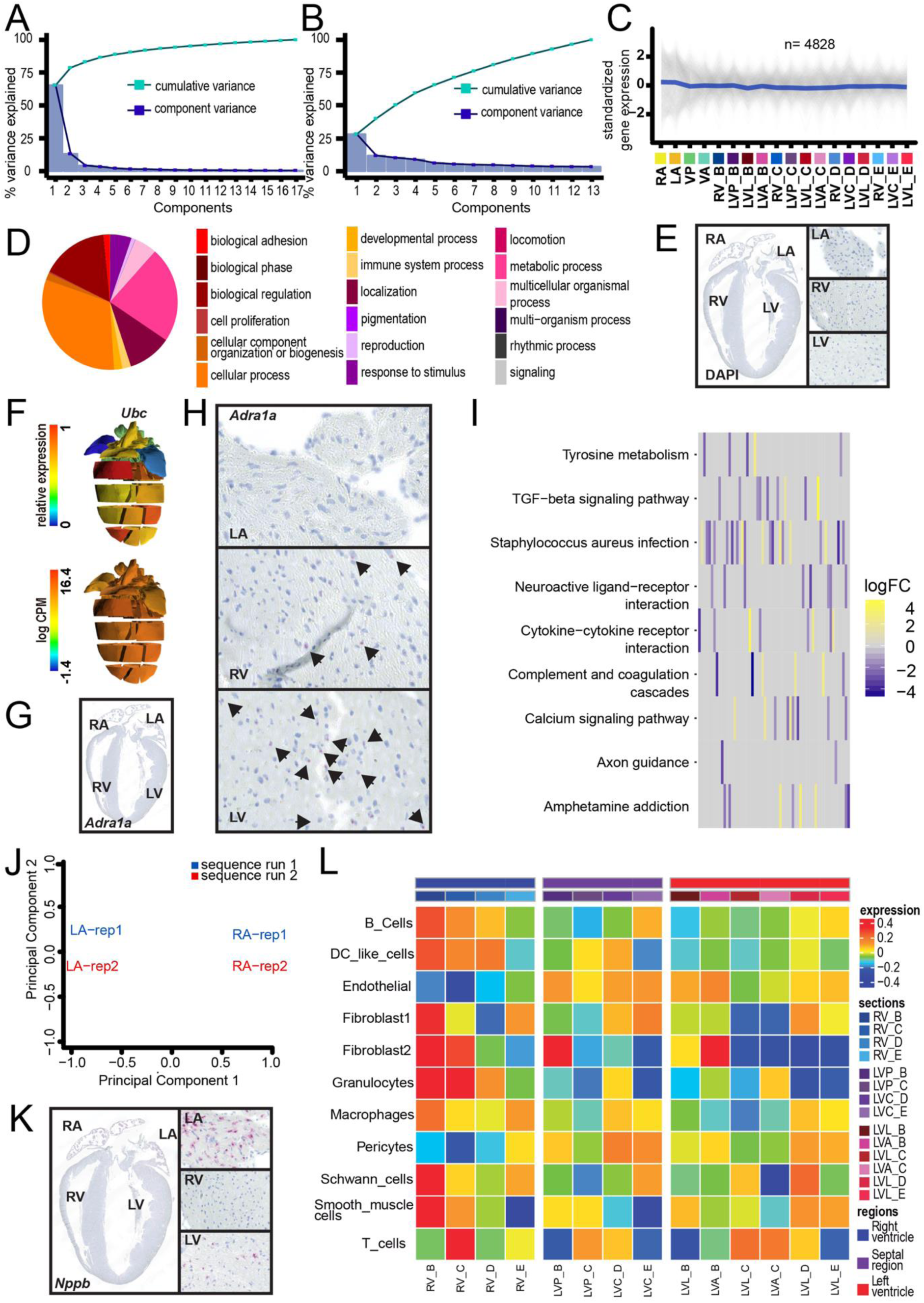
A-C) Variance explained by individual components, and cumulative variance across all components of correspondence analysis with data of all sections in cpm A), and ventricular sections in cpm B). C) All genes which had an alpha < 0.7 in soft cluster analysis and D) the proportions of these genes in enriched biological processes. E) RNA-scope DAPI control across main cardiac sections at 2.5X (left) and 10X (right) magnification. F) Relative and log CPM expression of *Ubc*. *Adra1a* expression at 2.5X magnification G), and 10X magnification H) across all cardiac sections. I) KEGG pathway enrichment heatmap with differentially expressed genes between the left and right atria. J) PCA of atrial sections and genes found in top enriched molecular functions of DGE analysis between the left and right atria. K) RNA-scope of *Nppb* expression across main cardiac sections at 2.5X (left) and 10X (right) magnification. L) Heatmap of average cell type expression across ventricular sections.

